# Simultaneous Inhibition of *ACLY* and *OGDH* Has a Synergistic Effect in Hepatocellular Carcinoma Cell Lines

**DOI:** 10.64898/2026.04.19.716936

**Authors:** Mehdi Dehghan Manshadi, Payam Setoodeh, Habil Zare

## Abstract

**Purpose:** Hepatocellular carcinoma (HCC) remains a leading cause of cancer-related mortality worldwide, and current therapies provide limited benefit without offering a definitive cure. This highlights the need for more effective therapeutic strategies. In this study, we investigated whether co-targeting the metabolic enzymes ATP citrate lyase (*ACLY*) and oxoglutarate dehydrogenase (*OGDH*) could induce synthetic lethality in HCC cells. Because valproic acid (VPA) and bempedoic acid (BA), inhibitors of *OGDH* and *ACLY*, have not previously been evaluated in combination in HCC, we investigated their potential to induce synthetic lethality in HCC cells.

**Methods:** Synthetic lethality was evaluated in HCC cell lines (Hep3B and Huh7) using pharmacological inhibition of *OGDH* and *ACLY* with VPA and BA, respectively. The effects of single-agent and combination treatments on cell proliferation were compared, and cytotoxicity was also assessed in the non-cancerous liver cell line THLE-2 to examine selectivity. Drug interaction effects were assessed based on the observed synergistic suppression of HCC cell growth relative to either treatment alone.

**Results:** Combined treatment with VPA and BA produced a strong synergistic effect in suppressing proliferation in Hep3B and Huh7 cells compared with either drug alone. In contrast, the combination caused little additional cytotoxicity in THLE-2 cells, suggesting a degree of cancer selectivity. These findings are consistent with previous reports identifying *USP13* as a metabolic regulator of *ACLY* and *OGDH* in multiple cancers, supporting the idea that the antiproliferative effects of *USP13* inhibition in HCC may be mediated through downstream effects on these two enzymes.

**Conclusions:** Co-targeting *ACLY* and *OGDH* shows promise as a synthetic lethal strategy in HCC and may provide a selective therapeutic approach with limited toxicity in non-cancerous liver cells. These findings support further investigation of *ACLY* and *OGDH* as a synthetic lethal pair and of combined VPA-BA treatment as a potential therapeutic strategy for HCC.

## Introduction

Synthetic lethality describes a genetic interaction in which the combined impairment of two or more genes results in cell death, whereas impairment of each gene individually is tolerated (1–3). This concept has become increasingly important in cancer research, providing a framework for the discovery of new therapeutics (4) and a powerful basis for cancer drug target identification (5). In particular, synthetic lethality offers a strategy for indirectly targeting proteins that have historically been difficult to address through conventional drug development, such as tumor suppressors with loss-of-function mutations and oncogenes activated by amplification or other genomic alterations. Despite these advantages, synthetic lethality remains an emerging therapeutic approach (6). Computational strategies, including systems biology, network-based analysis, machine learning, and genome-scale metabolic modeling, have been widely used to identify synthetic lethal gene sets for targeted therapy (7–10). However, broader clinical application remains constrained by the difficulty of identifying reliable gene pairs and the need for rigorous experimental validation. Recent advances in high-throughput CRISPR-Cas9 screening technologies have substantially accelerated the discovery of synthetic lethal interactions and their associated predictive biomarkers(4). In addition to identifying novel drug targets, synthetic lethal sets can support a range of therapeutic strategies, including drug repurposing (11), personalized medicine (12), and rational combination therapy (13,14).

Using computational analysis, we previously identified *OGDH* and *ACLY* as a synthetic lethal pair in hepatocellular carcinoma (HCC) (10). In the present study, we evaluated the therapeutic potential of this pair in HCC. *OGDH* (oxoglutarate dehydrogenase) and *ACLY* (ATP citrate lyase) encode enzymes with important roles in cellular metabolism. *OGDH* is a key enzyme in the tricarboxylic acid (TCA) cycle, catalyzing the conversion of α-ketoglutarate to succinyl-CoA, and is essential for cellular energy production, metabolic regulation, and oxidative stress responses (15). *OGDH* has also been implicated in several cancers, including gastric cancer (16). *ACLY* is a cytosolic homotetrameric enzyme that catalyzes the ATP-dependent conversion of citrate and CoA to acetyl-CoA and oxaloacetate, thereby linking mitochondrial carbohydrate metabolism to cytosolic lipid biosynthesis (15). *ACLY* is likewise central to cancer metabolism, and numerous *ACLY* inhibitors have been developed as anticancer agents (17). However, this synthetic lethal pair has been investigated only rarely in cancer research. In ovarian cancer, for example, one study showed that simultaneous suppression of *OGDH* and *ACLY* through *USP13* targeting may induce synthetic lethality by disrupting metabolism (18).

Using the DrugBank database (19), we selected two drugs for pharmacological inhibition of this pair. To target *OGDH*, we chose valproic acid (VPA, DB00313 (20)), a well-established antiepileptic drug. To target *ACLY*, we used bempedoic acid (BA, DB11936 (21)), a cholesterol-lowering agent. Our results demonstrate a significant synergistic effect of VPA and BA in HCC, highlighting the promise of drug repurposing and combination therapy as a potential therapeutic strategy.

## Materials and Methods

### Reagents

Valproic acid sodium salt (Cat# P4543) was obtained from Sigma-Aldrich and dissolved in ddH_2_O to prepare a 100 mM stock solution. Working concentrations of 0.25, 0.5, 1, 2, and 5 mM (22) were prepared in RPMI-1640 culture medium before treatment. Bempedoic acid (Cat# T3625) was purchased from TargetMol and prepared as a 1 mM stock solution in DMSO. For experimental treatments, working concentrations of 5, 10, 25, 50, and 100 µM (23,24) were prepared in RPMI-1640 medium. Fresh aliquots of all solutions were prepared for each experiment to ensure consistency and reliability.

### Cell Culture

The immortalized human liver epithelial cell line THLE-2 and the human HCC cell line Hep3B were purchased from ATCC (Manassas, VA, USA). The human HCC cell line Huh7 was provided by Dr. Robert Lanford of the Texas Biomedical Research Institute (San Antonio, Texas). Cell line identity was confirmed by short tandem repeat profiling, and all experiments were conducted using mycoplasma-free cells. Hep3B and Huh7 cells were cultured in RPMI-1640 medium supplemented with 10% fetal bovine serum (FBS; Gibco, Cat# 10082147), 0.5% of a 0.5 g/mL D-glucose stock solution, 1% of a 1 mM sodium pyruvate solution, and 1% penicillin-streptomycin (10,000 µg/mL). THLE-2 cells were cultured using the BEGM BulletKit (CC-3170; Lonza), which requires a specialized coating medium consisting of RPMI-1640 without glutamine supplemented with 0.01 mg/mL bovine serum albumin, 0.03 mg/mL type I collagen from bovine skin, and 0.01 mg/mL fibronectin from human plasma. All cell lines were maintained at 37°C in a humidified incubator containing 5% CO_2_.

### WST-1 Cell Viability Assay

The WST-1 cell proliferation assay reagent (Cat# 05015944001) was purchased from Sigma-Aldrich. Cells were seeded at a density of 1,000 cells per well in 96-well plates 24 hours before treatment. The following day, cells were treated with valproic acid, bempedoic acid, or their combination for 72 or 96 hours. After treatment, WST-1 reagent was added to each well at a 1:10 ratio, and the plates were incubated for 60 minutes. Absorbance was measured at 440 nm, as shown in Figure 1A. Higher absorbance values indicate greater cell viability. Fold change was calculated as the ratio of the treatment value to the control value. A value below 1 indicates inhibition, whereas a value above 1 indicates activation or stimulation.

**Figure 1.**
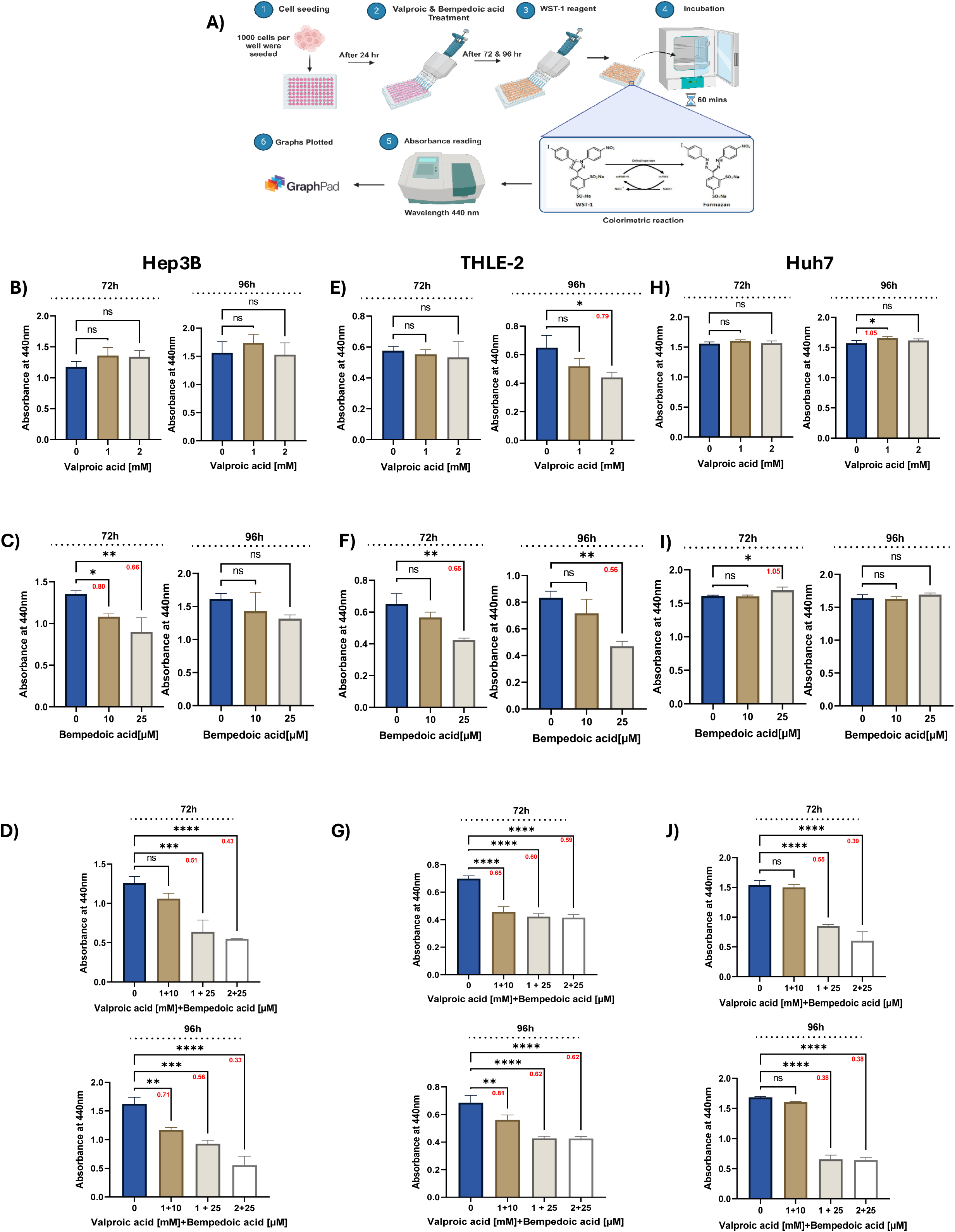
WST-1 cell viability assays were performed to evaluate the effects of VPA and BA, administered alone or in combination, on THLE-2, Hep3B, and Huh7 cells. (**A**) Schematic overview of the assay workflow. (**B–D**) Effects of VPA (0, 1, and 2 mM), BA (0, 10, and 25 μM), and their combinations, respectively, on THLE-2 cells. (**E–G**) Corresponding effects on Hep3B cells. (**H–J**) Corresponding effects on Huh7 cells. Cell viability was assessed 72 and 96 hours after treatment, as indicated at the top of each panel. Red-highlighted values represent fold-change values relative to the control; values <1 indicate inhibition, whereas values >1 indicate stimulation. *P < 0.05, **P < 0.01, ***P < 0.001, ****P < 0.0001, ns, not significant. Statistical analysis was performed using one-way ANOVA. Data are presented as mean ± SD.

### Statistical analysis

Data were analyzed using R Version 4.4.3 (25), and statistical differences were assessed by one-way analysis of variance (ANOVA) followed by Dunnett’s multiple-comparisons test (26). Results are presented as mean ± standard deviation (SD), and statistical significance was defined as *P* < 0.05. Graphs were generated using GraphPad Prism version 10.4.1 (27).

## Results

In our previous study(10), we conducted a comprehensive synthetic lethality analysis in HCC using Rapid-SL(28) and rFASTCORMICS(29), which identified 282 SL and higher-order SL sets as potential drug targets. These targets were selected based on their predicted ability to impair HCC tissue while minimizing adverse effects on the liver and 12 additional non-cancerous tissues.

To facilitate experimental validation and improve the feasibility of the present study, we focused on double SL pairs in which both genes were functionally linked within the same biological pathway. Applying this criterion reduced the candidate list to five double SL pairs: *OGDH*/*ACLY*, *DLST*/*ACLY*, *ALDH1L1*/*ALDH1L2*, *CP*/*FTMT*, and *SLC23A1*/*SLC23A2*. Among these, *OGDH* and *ACLY* were selected as the most promising pair. Both genes are closely associated with central carbon metabolism and the TCA cycle. This choice was supported by the availability of well-established drugs targeting these genes, providing a strong basis for both therapeutic development and experimental validation. Specifically, valproic acid was used as the *OGDH* inhibitor; this drug is widely prescribed for certain seizure disorders (30,31). Bempedoic acid, a clinically approved lipid-lowering drug used in patients with hypercholesterolemia (32), was selected as the *ACLY* inhibitor. Together, these agents provided a practical and readily available pharmacological strategy for targeting the selected synthetic lethal gene pair.

### Effects of VPA and BA on the Viability of THLE-2, Hep3B, and Huh7 Cells

The effect of VPA on Hep3B and Huh7 cells has been reported previously (33). In the present study, we evaluated its synergistic effect in combination with BA. To assess treatment specificity, THLE-2 cells were included as a non-cancerous control to determine whether this cell line showed increased sensitivity to the VPA-BA combination.

### Effects of Individual Drug Treatment

We first tested each drug individually to determine effective concentrations for combination treatment. VPA did not significantly reduce Hep3B cell viability except at the relatively high dose of 5 mM (Supplementary Figure 1B). Likewise, THLE-2 cell viability was not affected by VPA after 72 hours of treatment at any tested dose (Supplementary Fig. 1A). After 96 hours, THLE-2 viability was moderately reduced at 2 and 5 mM, with fold-change values of 0.79 and 0.67, respectively (*P* < 0.05; Supplementary Fig. 1A). In contrast, BA reduced the viability of both Hep3B and THLE-2 cells in a dose-dependent manner over the concentration range of 5-100 µM (Supplementary Fig. 1C, D).

Based on these individual treatment results, we selected 1 and 2 mM VPA for combination with 10 and 25 µM BA. These doses were chosen to minimize effects on non-cancerous cell viability. At the selected concentrations, VPA alone did not inhibit viability in Hep3B or THLE-2 cells, except for a moderate but statistically significant effect in THLE-2 cells treated with 2 mM VPA for 96 hours (Figure 1B and 1E). Likewise, BA alone produced either no reduction or only a moderate reduction in viability in the two cell lines, with fold-change values remaining at or above 0.56 (Figure 1C and 1F).

### Combination of Two Drugs

We next quantified the effects of combined treatment across different dose combinations (Figure 1D, 1G, and 1J). In Hep3B cells, the combined treatment produced a pronounced synergistic effect, suggesting enhanced therapeutic efficacy (Figure 1G). This effect was reflected by a marked reduction in viability relative to the corresponding single-drug treatments. Specifically, after 72 hours, treatment with 1 mM VPA plus 25 μM BA reduced Hep3B cell viability to a fold-change value of 0.51, compared with 1.14 and 0.80 for VPA and BA alone, respectively (Figs. 1E-G). A similar pattern was observed with 2 mM VPA combined with 25 μM BA after 72 hours, yielding a fold-change value of 0.43, compared with a non-significant change for VPA alone and 0.66 for BA alone. At lower-dose combinations, a longer treatment duration was required before a clear inhibitory effect became evident. For example, 1 mM VPA combined with 10 μM BA did not significantly reduce Hep3B viability after 72 hours (fold change: 0.85, compared with 0.80 and 1.16 for the individual treatments), but the same combination significantly reduced viability after 96 hours (0.72, compared with 1.13 and 0.89, respectively). Overall, the effects of combination treatment were stronger after 96 hours than after 72 hours. Notably, dose combinations that produced no significant inhibition when used individually led to significant inhibition when used together. The strongest combinatorial effect was observed after 96 hours with 2 mM VPA plus 25 μM BA, which reduced the fold-change value to 0.33, whereas each drug alone caused only non-significant inhibition.

After observing this synergistic effect in Hep3B cells, we evaluated the same combinations in the non-cancerous THLE-2 cell line. In all three tested combinatorial conditions, THLE-2 viability was significantly reduced after both 72 and 96 hours (Figure 2D). Interestingly, the inhibitory effect in THLE-2 cells appeared to diminish as the dose increased, suggesting a relative tolerance of non-cancerous cells to the combined treatment at higher concentrations. Consistent with this observation, THLE-2 cells were less sensitive than Hep3B cells at the higher dose combinations. To quantify this difference, we calculated the percentage change relative to the control and compared the inhibitory effects of the combined treatments across cell lines (Figure 2).

**Figure 2.**
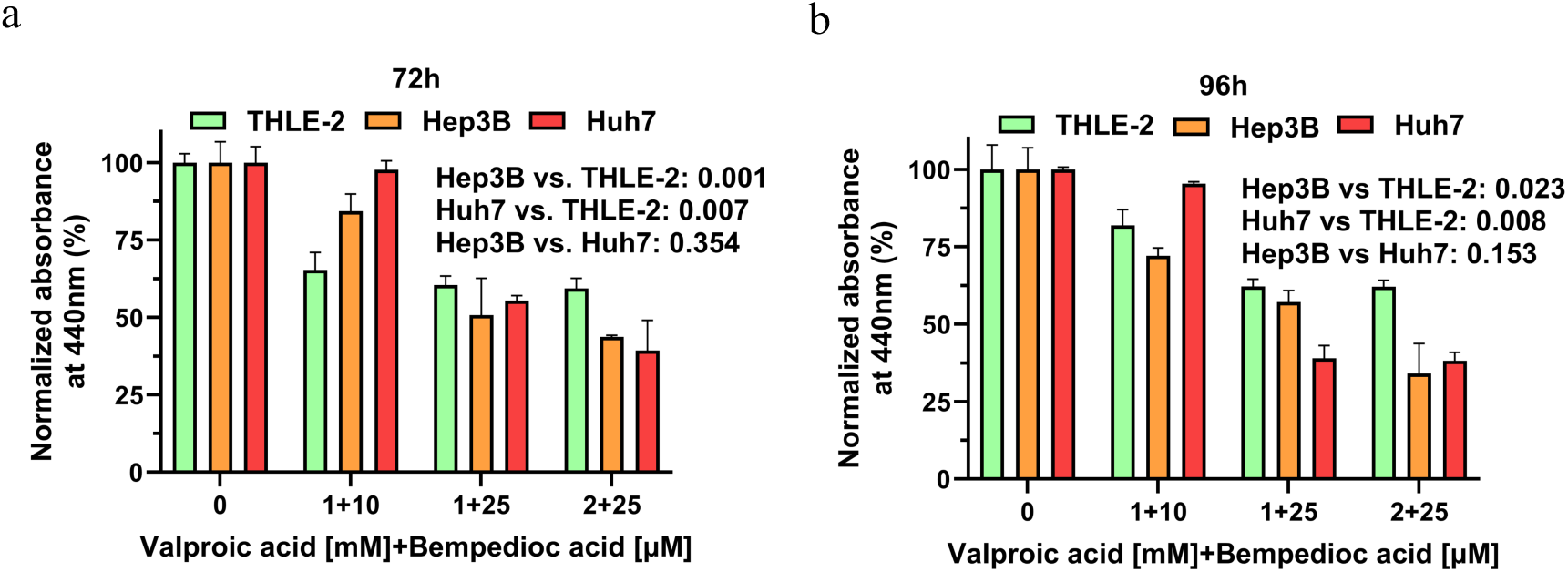
Comparison of the inhibitory effects of combined VPA and BA treatment across different cell lines after 72 hours (**a**) and 96 hours (**b**). The y-axis shows fold-change inhibition relative to the control for each cell line. Data are presented as mean ± SD.

The difference between the cancerous and non-cancerous cell lines was most evident with 2 mM VPA plus 25 μM BA. Under this condition, THLE-2 viability remained at 63% after 72 hours and 64% after 96 hours, whereas Hep3B viability was markedly lower at 43% and 41%, respectively. These findings indicate that the combined treatment exerted a significantly greater effect on Hep3B cells than on THLE-2 cells. To confirm the dose-dependent synergistic effect of the two drugs in HCC cells, we applied the same treatment protocol to a second human HCC cell line, Huh7. Neither drug exhibited inhibitory activity in Huh7 cells when used alone (Figs. 1H and 1I). In contrast, combined treatment produced a clear dose-dependent synergistic inhibitory effect, with fold-change values ranging from 0.55 to 0.38 (Figure 1J). Taken together, these findings suggest that the observed synthetic lethal effect is relatively selective for HCC cells, with only limited synergistic activity in non-cancerous liver cells, thereby supporting its therapeutic potential.

## Discussion

In this study, we evaluated the effect of simultaneously targeting *OGDH* and *ACLY* as a synthetic lethal pair in HCC cells. VPA and BA were selected as pharmacological inhibitors of *OGDH* and *ACLY*, respectively. Our results demonstrated a strong synergistic effect when these two drugs were combined at appropriate doses in the Hep3B and Huh7 HCC cell lines, compared with treatment using either drug alone. To assess selectivity, we also tested the same combination in the non-cancerous liver cell line THLE-2. We found that higher-dose combinations significantly impaired the viability of cancer cells while producing a substantially weaker synergistic effect in THLE-2 cells. These results therefore support the hypothesis that *ACLY* and *OGDH* may function as a synthetic lethal pair in HCC and that their co-inhibition may induce a selective synthetic lethal effect in HCC cells.

Our findings are consistent with previous studies implicating *USP13* in cancer metabolism. *USP13* has been identified as a driver of ovarian cancer metabolism, and its inhibition has been reported to suppress ovarian tumor progression (18,34). Because *USP13* regulates both *ACLY* and *OGDH*, earlier studies have suggested that *USP13* targeting may induce metabolic synthetic lethality through coordinated downregulation of these two genes in ovarian cancer. This hypothesis is further supported by reports linking *USP13* to *ACLY* and *OGDH* in multiple cancers (35) and by evidence of *USP13* upregulation in HCC (36–38). Given the broad cellular functions of *USP13*, its inhibition may be associated with off-target effects. In contrast, our results suggest that direct co-inhibition of *ACLY* and *OGDH* may capture an important component of the therapeutic effect of *USP13* targeting while offering a potentially more selective and safer strategy. In addition, our study provides preliminary dosing guidance for a potential VPA-BA combination approach in HCC, which may inform future preclinical studies and validation in human organoid models.

Both VPA and BA have previously been investigated individually in relation to HCC or liver biology. VPA has been reported as a potential strategy for overcoming sorafenib resistance in HCC (39,40), and has also been shown to significantly reduce the viability of PLC/PRF/5 cells (41). Furthermore, previous work has suggested that combining VPA with other therapeutic agents may represent a promising treatment strategy for HCC (42). Although the direct effect of BA on HCC has not, to our knowledge, been specifically investigated, BA has been shown to reduce hepatic steatosis and fibrosis in mouse models, suggesting possible benefits in liver disease (43,44). Taken together, the existing literature supports the biological plausibility of our findings. It highlights the potential of the VPA-BA combination treatment in HCC, warranting further investigation of its synergistic effects and therapeutic value.

A key limitation of this study is the use of a non-cancerous cell line to assess the selectivity of the identified synthetic lethal interaction. Although this model provides an initial indication of selectivity, it does not fully reflect the complexity of normal liver tissue and retains cell-line-specific characteristics. Future studies using more physiologically relevant models, such as primary hepatocytes or liver organoids, will be necessary to better define the specificity and translational relevance of this proposed synthetic lethal interaction. In addition, whereas BA selectively inhibits *ACLY* (*45*), VPA is a histone deacetylase inhibitor with broad effects extending beyond *OGDH* (*46*). Future work could therefore use CRISPR-Cas9 (47) or siRNA(48)-based approaches to more precisely dissect the individual contributions of *ACLY* and *OGDH* to HCC progression and to validate their functional roles.

## Conclusion

Our findings highlight the therapeutic promise of co-targeting *ACLY* and *OGDH* as a synthetic lethal pair in hepatocellular carcinoma. Combined treatment with VPA and BA produced a synergistic inhibitory effect on cancer cell proliferation while exerting a comparatively smaller effect on non-cancerous cells, suggesting a degree of selectivity that may be advantageous for cancer therapy.

## Acknowledgements

We acknowledge Dr. Nagesh K. Panchal for performing the cell viability experiments and analyzing the resulting data. We thank Dr. Robert Lanford for providing the Huh7 cell line. The authors acknowledge assistance from an AI-based language tool (ChatGPT, OpenAI) used solely for language refinement.

## Disclosure

The authors report no conflicts of interest in this work. This paper has been uploaded to BioRxiv as a preprint: https://www.biorxiv.org/content/10.64898/2026.04.19.716936v1.

## Notes

### Competing Interest Statement

The authors have declared no competing interest.

### Summary of Updates

Dr. Sun Luzhe and Dr. Nagesh K. Panchal have been removed from the author list at their request. Dr. Panchal's contribution to the study has been included in the Acknowledgments section.

### Funding

- NIA: R01AG057896, 1RF1AG063507, R01AG068293, 6 1R01AG0665241A, 1R01AG065301, P30 AG066546, and 1R21AG087907
- NINDS: RF1NS112391, U19NS115388, and R015R01NS017950-39

