## Supplementary Figures for "Simultaneous Inhibition of *ACLY* and *OGDH* Has a Synergistic Effect in Hepatocellular Carcinoma Cell Lines"

Mehdi Dehghan Manshadi, Nagesh Kishan Panchal, Lu-Zhe Sun, Payam Setoodeh, Habil Zare

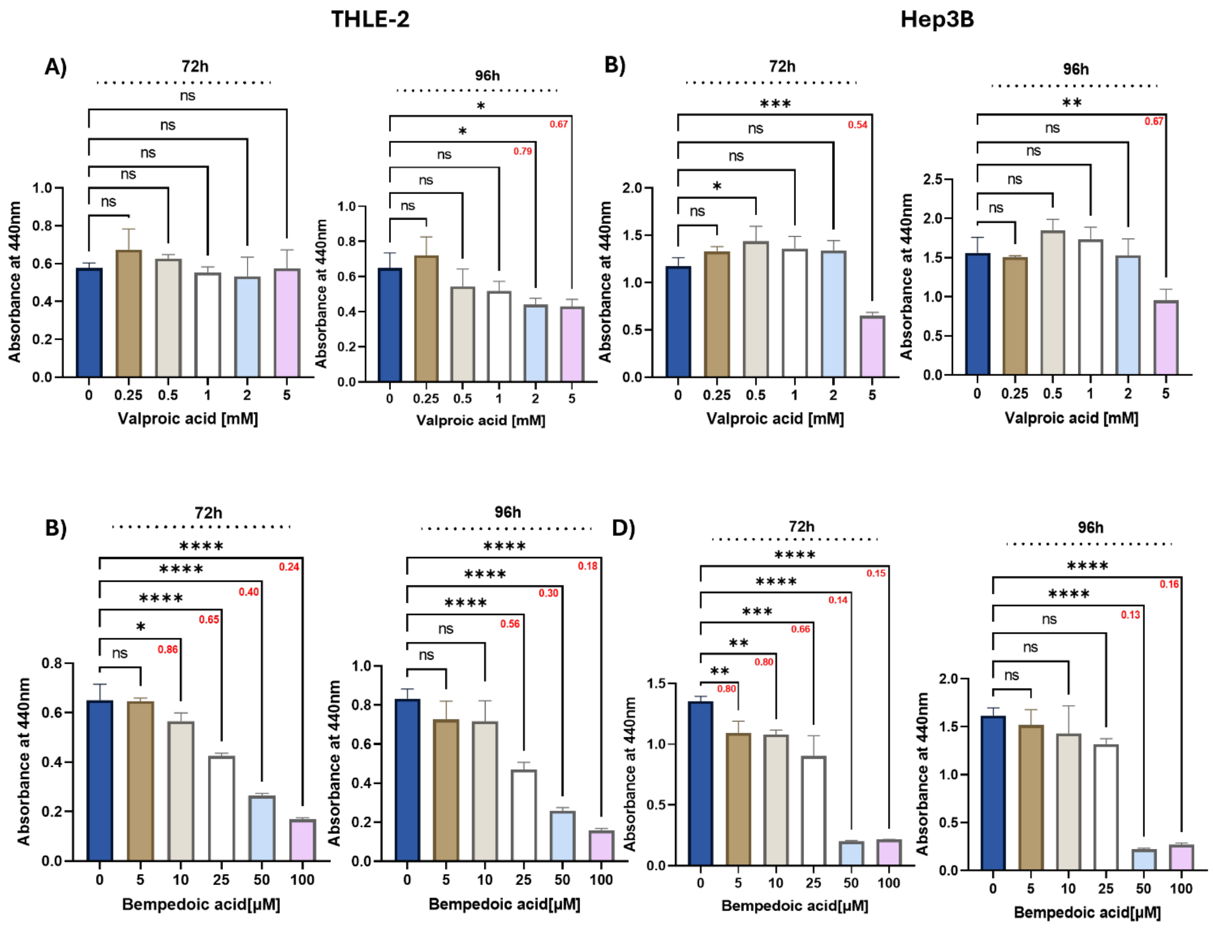


**Supplementary Figure 1.** The figure demonstrates the results of cell viability assays conducted using the WST-1 to assess the effects of valproic acid and bempedoic acid on the viability of THLE-2 and Hep3B cells. **A & C)** display the effects of valproic acid (0, 0.25, 0.5,1, 2, and 5 mM), and **B & D)** display the effects of bempedoic acid (0, 5,10, 25, 50, and100 μM) on THLE-2 and Hep3B cells, respectively. Cell viability was measured at 72-hour and 96-hour post-treatment as shown. Red highlighted values denote fold change inhibitory effects relative to the control. Values greater than one represent stimulation, whereas values less than one indicate inhibition. **P < 0.05, **P < 0.01, ***P < 0.001, ****P < 0.0001, ns- non-significant*. Analysis was performed using one way ANOVA. Data are shown as mean ± SD.
